# Characterization of new highly selective pyrazolo[4,3-d]pyrimidine inhibitor of CDK7

**DOI:** 10.1101/2023.02.24.529844

**Authors:** Markéta Kovalová, Libor Havlíček, Stefan Djukic, Jana Škerlová, Miroslav Peřina, Tomáš Pospíšil, Eva Řezníčková, Pavlína Řezáčová, Radek Jorda, Vladimír Kryštof

## Abstract

Targeting cyclin-dependent kinase 7 (CDK7) provides an interesting therapeutic option in cancer therapy because this kinase participates in regulating the cell cycle and transcription. Here, we describe a new trisubstituted pyrazolo[4,3-*d*]pyrimidine derivative, LGR6768, that inhibits CDK7 in the nanomolar range and displays favourable selectivity across the CDK family. We determined the structure of fully active CDK2/cyclin A2 in complex with LGR6768 at 2.6 Å resolution using X-ray crystallography, revealing conserved interactions within the active site. Structural analysis and comparison with LGR6768 docked to CDK7 provides an explanation of the observed biochemical selectivity, which is linked to a conformational difference in the biphenyl moiety. In cellular experiments, LGR6768 affected regulation of the cell cycle and transcription by inhibiting the phosphorylation of cell cycle CDKs and the carboxy-terminal domain of RNA polymerase II, respectively. LGR6768 limited the proliferation of several leukaemia cell lines, triggered significant changes in protein and mRNA levels related to CDK7 inhibition and induced apoptosis in dose- and time-dependent experiments. Our work supports previous findings and provides further information for the development of selective CDK7 inhibitors.

**Graphical abstract:** 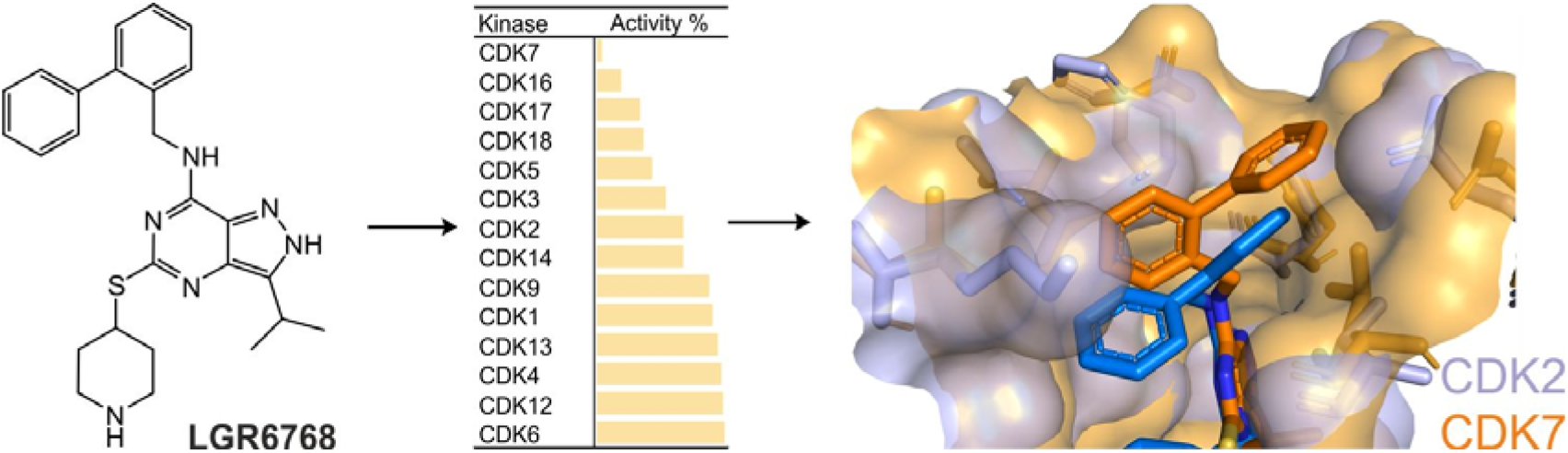

## 1. Introduction

Cyclin-dependent kinase 7 (CDK7) is a serine-threonine kinase that plays a crucial role in regulating the cell cycle and transcription, linking these processes together. CDK7 forms a trimeric complex with cyclin H and MAT1, which is called a CDK-activating complex (CAK), that is responsible for activating CDK1 and CDK2 by phosphorylating their T-loops at T160 and T161, respectively. Moreover, CDK7 regulates transcription as a part of the general transcription factor TFIIH, which is recruited to a Mediator–preinitiation complex responsible for phosphorylating the carboxy-terminal domain of RNA polymerase II at serine 5, which facilitates the initiation of transcription [1].

While cell cycle CDKs or cyclins are often amplified or overexpressed in cancer, the involvement of transcriptional CDKs (tCDK) in the initiation and development of cancer is unclear. It was proposed that CDK7 and other tCDKs are necessary for the expression of highly expressed antiapoptotic genes, genes controlled by oncogenic transcription factors and fusion chimaeras (e.g., MYC, MLL1-AF9), and genes associated with cancer-specific super-enhancer elements. It was recently discovered that the genes encoding tCDKs in cancers are usually characterized by a loss in copy numbers. Among others, CDK7 showed the most significant copy-number loss across more than 10,000 assessed tumours [2]. These genetic alterations are correlated with increased sensitivity of cancer cells to DNA-damaging agents, which is mechanistically linked to the repression of genes encoding for the DNA damage response pathways. Due its essential roles in cell proliferation and transcription, CDK7 has become intensively studied as a possible target in cancer treatment.

In the last decade, several selective CDK7 inhibitors have been developed, which function either as traditional ATP competitors or as irreversible inhibitors. Currently, six compounds are in clinical studies (Figure 1). BS-181 was described as the first noncovalent selective CDK7 inhibitor [3] built on a pyrazolo[1,5-*a*]pyrimidine scaffold. Preclinical studies have shown its antitumour activity on several tumour types, but BS-181 exerts poor drug-like properties [4-6]. By modifying BS-181, the clinical candidate samuraciclib (also known as ICEC0942 or CT7001) was produced with improved selectivity towards CDK7 and strong antitumour effects in breast and colorectal cancer xenografts [7]. Further analogues of BS-181, LDC4297 and LDC3140, were built on the pyrazolo[1,5-*a*][1,3,5]triazine scaffold and displayed high specificity to CDK7 and selectivity [8]. Recently, indolylpyrimidine SY-5609 was reported as a subnanomolar, highly selective, noncovalent and orally available inhibitor of CDK7 with favourable ADME properties and is currently being evaluated in phase I clinical trials [9].

**Figure 1.**
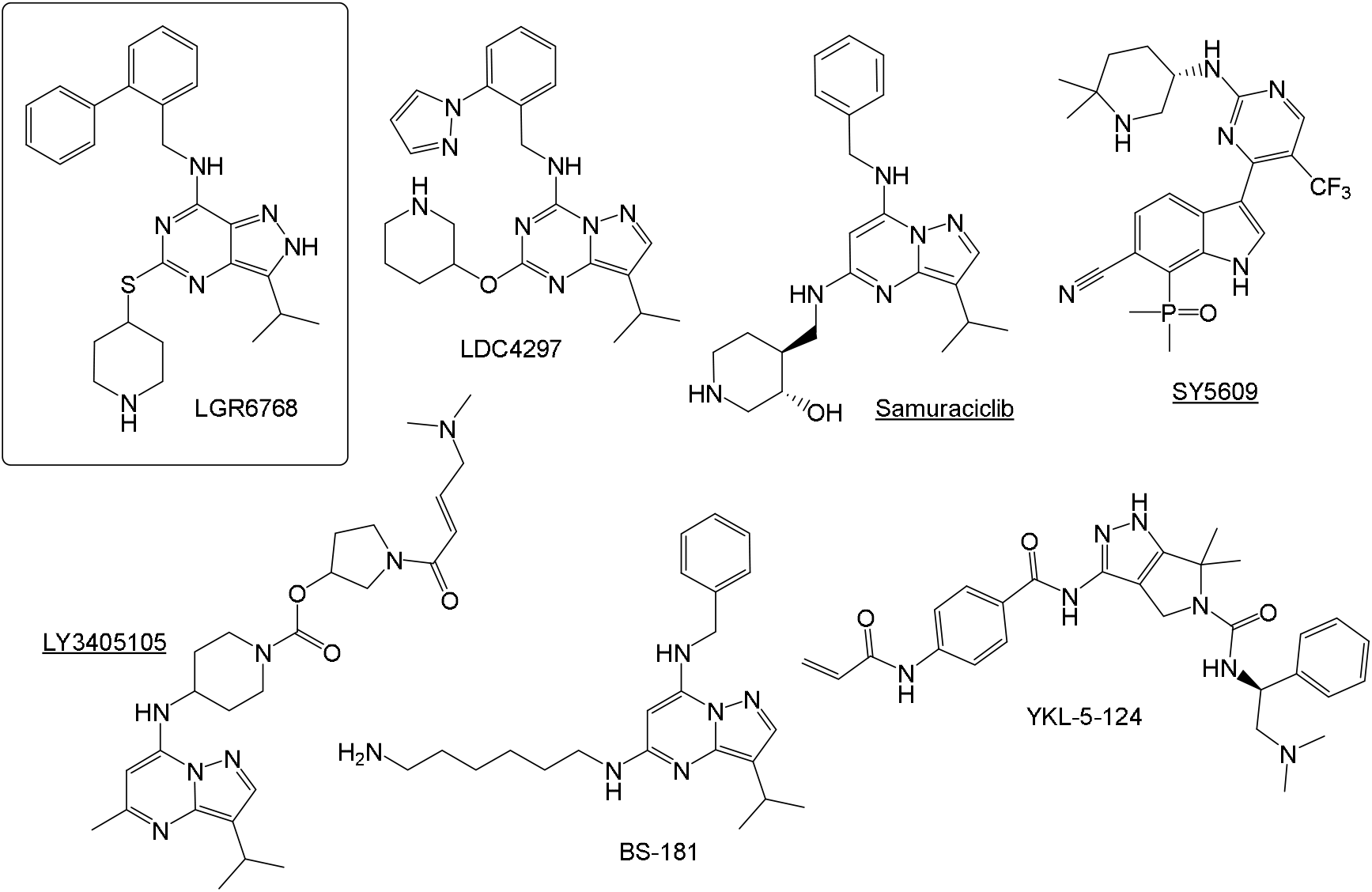
Structure of LGR6768 and some known CDK7 inhibitors (the underlined inhibitors were registered in clinical trials).

Selective CDK7 inhibition can be achieved by targeting reactive Cys312 residues outside the ATP pocket. THZ1 was described as the first covalent nanomolar CDK7 inhibitor [10] and was tested on many cancer models [11-15]. Unfortunately, THZ1 exhibited short plasma stability, but its position isomer THZ2 showed improved pharmacokinetics and antitumour activities [16]. Subsequent development led to the discovery of structurally related SY-1365 (mevociclib) [17], the first CDK7 covalent inhibitor that entered clinical trials. Its antitumour properties were observed in several xenograft models, either as monotherapy or in combination with venetoclax [17]. The combination of the THZ1 warhead and core from the PAK4 inhibitor PF-3758309 also led to the development of YKL-5-124 [18]. Although YKL-5-124 showed potency similar to that of THZ1, the compound exhibited a different phenotype in cancer cells and did not change global RNA polymerase II phosphorylation, probably due to its kinase selectivity profile.

Recently, other CDK7 inhibitors have been introduced, some of which have already entered clinical trials, namely, LY3405105, Q901 and XL102. Nevertheless, limited information about their mode of action, kinase selectivity or structure has been disclosed [19-23].

We have previously reported several series of potent 3,5,7-trisubstitued pyrazolo[4,3-*d*]pyrimidine CDK inhibitors with strong anticancer activity in vitro and in vivo [24-27]. Recently, we also showed the ability of some pyrazolo[4,3-*d*]pyrimidines to act as molecular glue, causing rapid and selective proteasome-dependent degradation of cyclin K [28]. In the current study, we describe that the change of the 4-phenylbenzyl ring to a 2-phenylbenzyl at the 7 position of pyrazolo[4,3-*d*]pyrimidine confers a significant change in the desired selectivity for CDK7. Here, we introduce the novel CDK7 inhibitor LGR6768, confirm its preference for CDK7 and describe its antiproliferative effect on several leukaemia cell lines.

## 2. Results and discussion

### 2.1. Compound design and structural rationale

The design of LGR6768 (5-(piperidin-4-yl)thio-3-isopropyl-7-(2-phenylbenzyl)amino-1(2)H-pyrazolo[4,3-*d*]pyrimidine) was inspired by the binding of previously described 3,5,7-trisubstituted pyrazolo[4,3-*d*]pyrimidines in the CDK2/cyclin cavity (PDB: 3PJ8, 6GWA, 7QHL) [24,25,28]. Conserved isopropyl at position 3 was retained, whereas incorporation of the piperidine-thio chain at position 5 and the biphenyl moiety at position 7 partly reflected the structures of known CDK7 inhibitors [8,29]. Compound LGR6768 was then synthesized as described previously by two subsequent reactions from pyrazolo[4,3-*d*]pyrimidine-5,7-dithiol [25,27]; for details, see Figure S1.

We first determined the crystal structure of fully active Thr160-phosphorylated CDK2 with cyclin A2 (residues 175-432) in complex with LGR6768 at 2.6 Å resolution (PDB: 8B54) to confirm the orientation in the CDK2 active site cavity. Data collection and refinement statistics are listed in Table S1, and the electron density maps are shown in Figure S2. The pyrazolo[4,3-*d*]pyrimidine core of the inhibitor forms direct hydrogen bonds with Glu81 and Leu83 residues. In addition, water-mediated hydrogen bonds are formed between N4 of the inhibitor core and the Lys33, Glu51, and Asn132 side chains, as well as the main-chain nitrogen of Asp145. The thioether-linked piperidine moiety adopts a twisted-boat conformation that is stabilized by a hydrogen bond with Gln131, together with additional water-mediated interactions with the Asp86 carboxyl. The biphenyl moiety is oriented towards the active site opening, forming hydrophobic interactions with Ile10, Asp86, Lys89 and Phe82 (Figure 2B, Figure S3). Finally, the isopropyl substituent is buried into the hydrophobic pocket, interacting with Val64, Leu134, Ala144, and the gatekeeper Phe80 (Figure S3).

**Figure 2.**
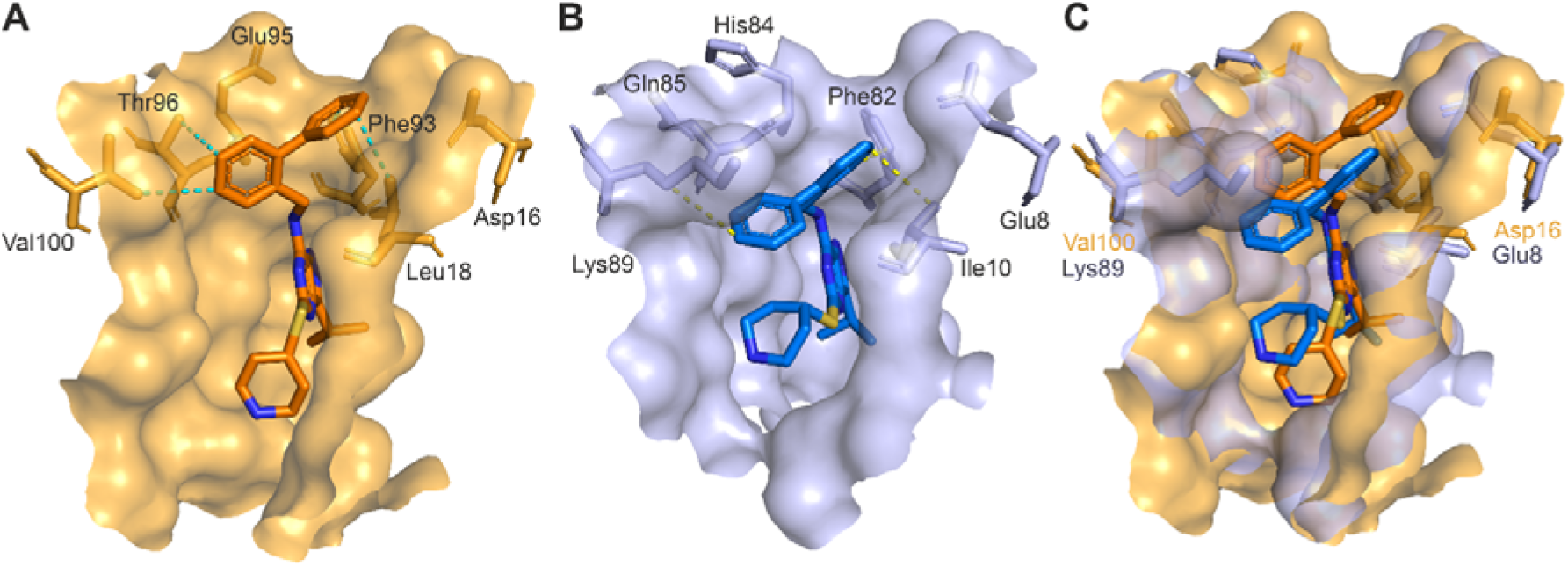
LGR6768 binding poses in the active sites of CDK7 and CDK2. (A) LGR6768 docked into the CDK7 cryo-EM structure (PDB: 7B5O). (B) CDK2 cocrystal structure with LGR6768 (PDB: 8B54). (C) Alignment of both binding poses and cavities of CDK7 and CDK2 shows structural differences in CDK7 (orange) and CDK2 (blue). Selected protein residues forming the edge of the cavity are shown as sticks. Hydrophobic interactions of the biphenyl moiety of LGR6768 with protein residues are shown as dashed lines. Ligand heteroatoms are blue (nitrogens) and yellow (sulfur).

Next, molecular docking of LGR6768 into the cryo-EM structure of CDK7 (PDB: 7B5O) was performed to determine its selectivity for CDK7. The best binding pose of the inhibitor in the active site of CDK7 (ΔG_Vina_ = -9.4 kcal/mol) is very similar to that in CDK2 (Figure 2A, B); the pyrazolo[4,3-*d*]pyrimidine core forms direct hydrogen bonds with Asp92 and Met94. However, key differences were observed when comparing the binding of LGR6768 in the active cavity of CDK2 and CDK7. Generally, the active site cavity of CDK7 is much wider and more open to a possible interaction (Figure 2C). This could be the reason why LGR6768 is in a more relaxed conformation in CDK7 than in CDK2 and why the piperidine moiety points deeper in the cavity to interact with Asp155. Next, the isopropyl group of LGR6768 protrudes into the analogical hydrophobic pocket in CDK7 interacting with Ala39, Ile75 and the gatekeeper Phe91 (Figure 2B, Figure S3). However, the isopropyl group is rotated by approx. 180° between the crystal structure in CDK2 and the modelled pose in CDK7, similar to inhibitor BS-181 in CDK7 [3]. The most evident difference in the binding pose of LGR6768 is the switched/flipped orientation of the biphenyl moiety, which points towards the active site opening in the CDK2 crystal structure but is oriented to the other side of the opening in the modelled pose in CDK7, pointing to the surface at the very edge of the active site cavity. Structural differences between CDK2 and CDK7 at the edge of the binding cavity could explain the opposite orientations of the biphenyl substituent. First, the cavity opening widens due to the substitution of Lys89 in CDK2 by Val100 in CDK7. More importantly, residue Glu8 in CDK2 forms a hump on the surface of the cavity edge; this observation clashes with the biphenyl moiety binding mode observed in CDK7, in which the biphenyl outer ring to point towards the surface due to homologous Asp16, which is oriented away from the cavity opening (Figure 2C). In addition, the substitution of Ile10 in CDK2 by Leu18 in CDK7 seems to be important as well, since one of the terminal methyl groups of Leu18 in CDK7 points towards the outer phenyl ring of LGR6768. The biphenyl moiety of LGR6768 forms extensive hydrophobic interactions with CDK7 residues Leu18, Phe93, Thr96 and Val100 (Figure 2A, Figure S3). Of note, the different orientations of the biphenyl moiety in CDK2 and CDK7 correspond well with the different orientations of the benzylamine substituent of samuraciclib observed in the same kinases. These were termed the “ring-up” and “ring-down” conformations, which are related by a rotation by ∼120°, and occurred because the structural differences between CDK2 and CDK7 were the same [30]. Based on the previous knowledge, we believe that the biphenyl substituent position and interaction are key for the potency and selectivity of LGR6768 towards CDK7.

### 2.2. Kinase selectivity

The selectivity profiling of LGR6768 was performed on a panel of 50 kinases, including known off-targets of other structurally related pyrazolo[4,3-*d*]pyrimidines [25,28] or isosteres, such as roscovitine or CR8 [31]. Given the high structural similarity within the CDK family, we also evaluated LGR6768 against 14 CDKs to assess its selectivity across them. The compound was applied at a 1 µM concentration, and the experiment confirmed selective reduction of the enzymatic activity of CDK7 to 4%. Only a few other kinases were significantly inhibited under the same settings, including CDK16 from the CDK family (Figure 3A) and CK1δ (both with 19% residual activity) (Figure S4). Dose-dependent measurement revealed potent inhibition of recombinant CDK7 at low nanomolar concentrations with IC_50_ = 20 nM, which was at least 12-fold lower than for other CDKs (Table S2). Finally, the selectivity profile of LGR6768 for selected CDKs was compared to that of other nanomolar CDK7 inhibitors, namely, THZ1, LDC4297 and samuraciclib (Figure 3B). Our results indicated that those inhibitors displayed a lower selectivity index than that of LGR6768, clearly confirming the improved kinase selectivity of our candidate.

**Figure 3.**
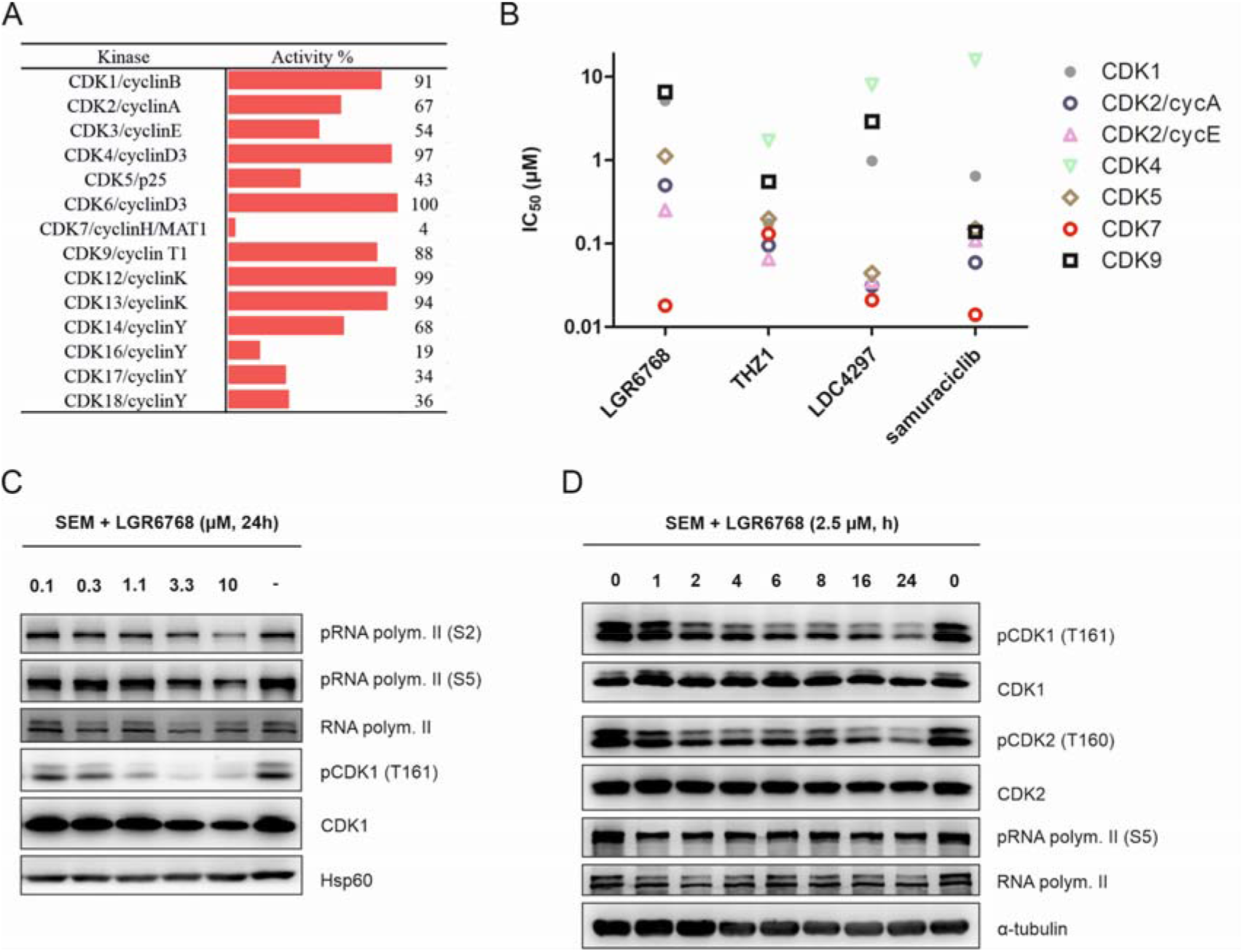
(A) CDK selectivity profile of LGR6768 assayed at a 1 µM concentration. (B) Comparison of kinase inhibition values of selected CDKs for LGR6768 and selected CDK7 inhibitors. IC_50_ values > 20 µM are not shown. (C, D) Immunoblotting analysis of CDK7-related proteins in the SEM cell line treated for the indicated times with LGR6768. Hsp60 and tubulin protein expression marked equal protein loading.

CDK7 is involved in regulating the cell cycle as CAK (CDK activating kinase) by activating phosphorylation of the T-loop of CDK1 and CDK2 at threonines 161 and 162, respectively. Immunoblotting analysis revealed that LGR6768 downregulates the phosphorylation of both CDKs in a dose- (Figure 3C) and time-dependent (Figure 3D) manner in the SEM cell line. Moreover, we also observed the expected dephosphorylation of RNA polymerase II (Figure 3C, D), although not as dramatic as was caused by the less selective CDK7 inhibitor THZ1 [10]. In fact, our results resemble those obtained with the CDK7 inhibitor YKL-5-124, which exhibited little effect on RNA polymerase II phosphorylation[18].

### 2.3. Antiproliferative and proapoptotic effects of LGR6768

Several CDK7 inhibitors showed antileukaemic effects in both AML and ALL cell lines [5, 10, 11, 17], and SY-1365 also showed antitumour activity in AML xenografts [17]. Therefore, we profiled LGR6768 on a panel of 10 cell lines derived from haematological malignancies (5 AML cell lines, 3 ALL cell lines, 1 lymphoma, 1 CML). The observed activities reached GI_50_ values between 0.44 and 4.4 µM except for K562 and CCRF-CEM cells (GI_50_ > 5 µM). Antiproliferative properties in seven selected cancer cell lines derived from different origins (breast, prostate, melanoma) showed higher GI_50_ values than those in AML and ALL cell lines, suggesting the antileukaemic potential of LGR6768 (Figure 4A).

**Figure 4.**
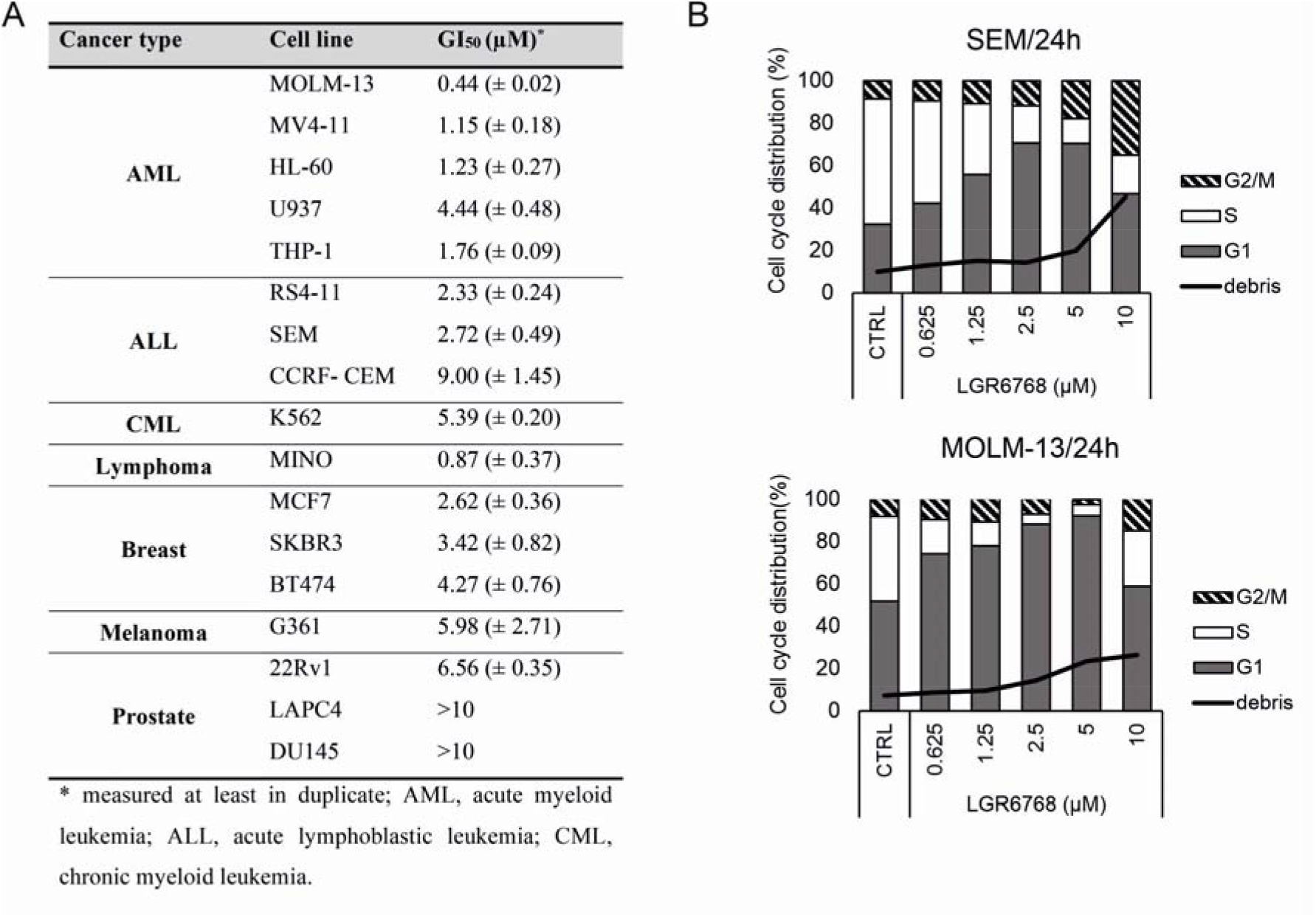
(A) Antiproliferative activity of LGR6768. (B) Effect of LGR6768 on cell cycle distribution in the SEM and MOLM-13 cell lines. Cells were treated for 24 hours with increasing concentrations of LGR6768 or vehicle.

The cell cycle phenotype caused by inhibition of CDK7 usually leads to a reduction in the number of S-phase cells and concomitant accumulation of cells in the G1 or G2/M phase of the cell cycle, which is specific for each cell line and the concentration of inhibitor [32]. One-day treatment with compound LGR6768 led to dose-dependent G1 arrest in both the SEM and MOLM-13 cell lines already observed at mid-nanomolar concentrations (Figure 4B). Significant apoptosis accompanied by G2/M arrested cells was observed solely at 10 µM. The effect of LGR6768 on the cell cycle of the most sensitive MOLM-13 cells showed a time-dependent tendency and deepened to low-nanomolar concentrations during the 3-day experiment (Figure S5).

Furthermore, we investigated the proapoptotic effect of LGR6768 in MV4-11, MOLM-13, and SEM cell lines. Cells were treated with increasing doses of LGR6768 for 24 hours and examined for apoptotic markers. As presented in Figure 5A, immunoblotting revealed only a moderate effect of LGR6768 on the tested cell lines, as documented by PARP-1 cleavage and detection of caspase-7 and caspase-9 fragments, mostly at 10 µM. Additionally, LGR6768 caused downregulation of the antiapoptotic protein Mcl-1 to a lesser extent, while no changes in Bcl-2 levels were detected, probably explaining its higher stability. In concordance with immunoblotting, a fluorometric caspase-3/7 activity assay confirmed the proapoptotic effect of LGR6768 in MOLM-13 and SEM cell lines after 24 hours of treatment (Figure 5B).

**Figure 5.**
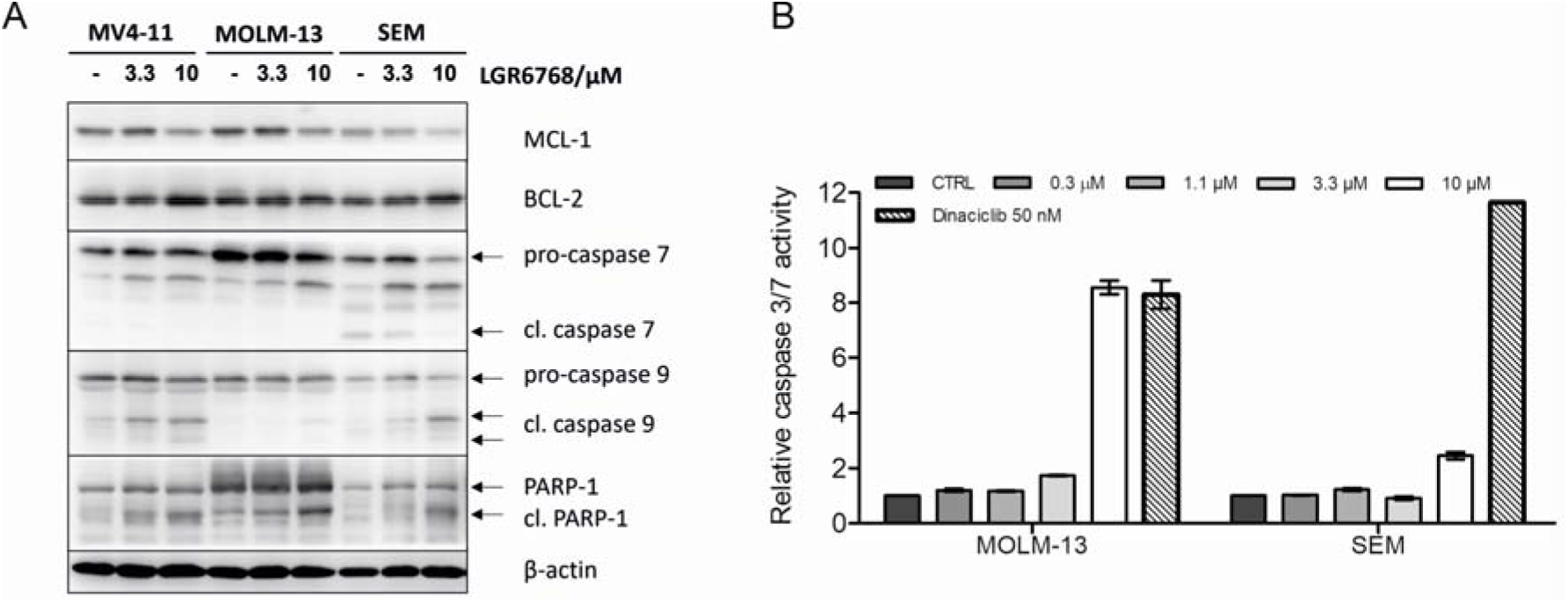
Proapoptotic effect of LGR6768. (A) Immunoblotting analysis of apoptosis-related proteins. MV4-11, MOLM-13 and SEM cells were treated for 24 hours with increasing concentrations of LGR6768. The actin level marked equal protein loading. (B) Activity of caspases 3/7 in MOLM-13 and SEM cell lines treated with LGR6768, dinaciclib or vehicle for 24 hours.

**Figure 6.**
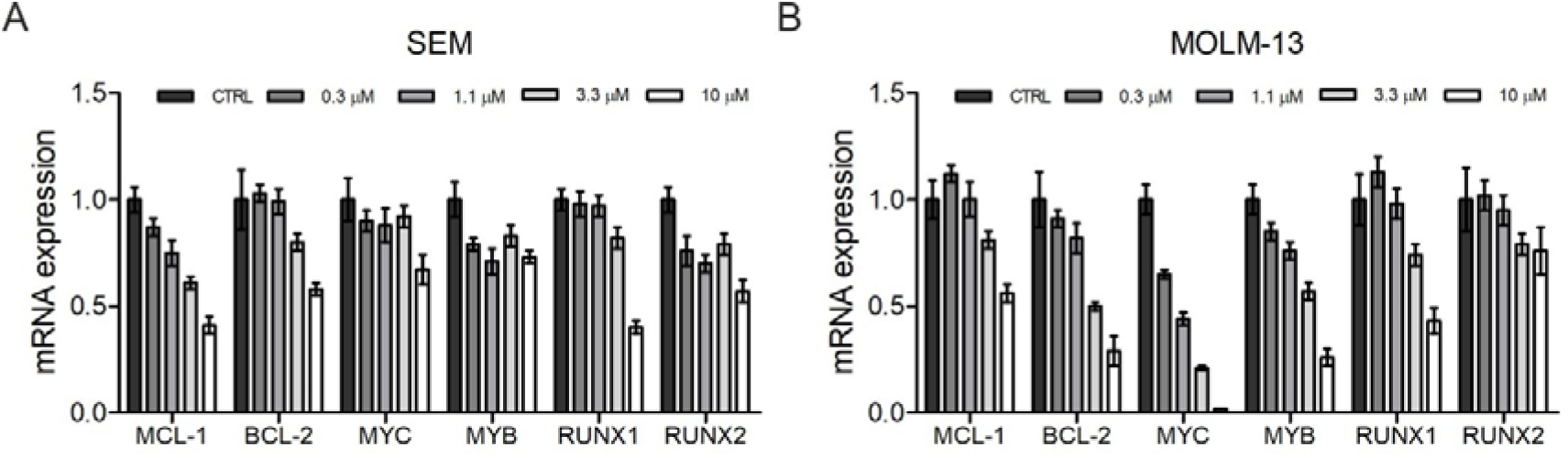
LGR6768 downregulates the expression of transcription factors and apoptosis-related genes in the SEM (A) and MOLM-13 (B) cell lines. Cells were treated with increasing concentrations of LGR6768 or vehicle for 4 hours. Relative expression levels were normalized to GAPDH and RPL13A genes (MOLM-13) or GAPDH and B2M genes (SEM).

### 2.4. Effect of LGR6768 on transcription targets

Furthermore, we investigated the inhibition of transcription in treated cells by analysing newly synthesized transcripts involved in apoptosis (e.g., Bcl-2, Mcl-1) or playing roles as oncogenic drivers in various haematological malignancies (e.g., RUNX, Myc, Myb). Those genes were previously established as genes sensitive to inhibition of global transcription and as fundamental genes important in targeting triple-negative breast cancer [16]. Their negative modulation by CDK7 inhibitors was highlighted in several studies, and as an example, targeting Jurkat ALL cells with low doses of THZ-1 was explained by diminishing the RUNX1-driven transcriptional program [16].

Transcriptional changes in mRNAs of anti-apoptotic genes *MCL-1* and *BCL-2* were observed to be suppressed with increasing doses of LGR6768 in SEM and MOLM-13 cell lines. These changes were examined after 4 hours of treatment and demonstrate the rapid action of the inhibitor and the ability to induce apoptosis. Most of the other transcripts were downregulated in a dose-dependent manner to 50% values. *MYC* expression decreased rapidly in MOLM-13 cells to < 1%, which could be explained by the highest sensitivity of MOLM-13 cells to LGR6768.

## 3. Conclusion

In recent years, CDK7 has emerged as an attractive target in cancer therapy due to its role in regulating both the cell cycle and transcription. Here, we describe the development of compound LGR6768, a new pyrazolo[4,3-*d*]pyrimidine derivative that potently and selectively inhibits CDK7. The selective targeting of CDK7 was confirmed by biochemical kinase profiling and supported by structural analysis. LGR6768 showed antileukaemic potential and downregulated the phosphorylation of CDK7 substrates, including CDK1, CDK2 and RNA polymerase CTD. Cell cycle analysis revealed a blockage in the G1 phase after treatment, and higher concentrations led to the induction of apoptosis accompanied by G2/M block, caspase activation, PARP-1 cleavage, and downregulation of the expression of antiapoptotic proteins at both the mRNA and protein levels. Moreover, LGR6768 downregulated the expression of several oncogenic drivers of haematological malignancies, indicating that targeting CDK7 is worth further exploration as a new therapeutic modality.

## 4. Experimental section

NMR spectra were recorded on a JEOL ECA-500 spectrometer operating at frequencies of 500.16 MHz (^1^H) and 125.76 MHz (^13^C) or on JEOL-ECZ 400/R spectrometer operating at frequencies of 399.78 MHz (^1^H) and 100.53 MHz (^13^C). ^1^H NMR and ^13^C NMR chemical shifts were referenced to the solvent signals; ^1^H: δ(residual CHCl_3_) = 7.25 ppm, δ(residual DMSO -*d*_5_) = 2.50 ppm; ^13^C: δ(CDCl_3_) = 77.23 ppm, δ(DMSO-*d*_6_) = 39.52 ppm. Chemical shifts are given on the δ scale [ppm], and coupling constants are given in Hz. Melting points were determined on a Kofler block and are uncorrected. Reagents were of analytical grade from standard commercial sources or were synthesized according to the referenced procedure. Thin layer chromatography (TLC) was carried out using aluminum sheets with silica gel F254 from Merck. Spots were visualized under UV light (254 nm). ESI mass spectra were determined using a Waters Micromass ZMD mass spectrometer (solution of sample in MeOH, direct inlet, coin voltage in the range of 10-20 V, trace amounts of HCOOH or NH_4_OH were used to influence ionization). Column chromatography was performed using Merck silica gel Kieselgel 60 (230-400 mesh). All compounds gave satisfactory elemental analyses (± 0.4%).

### 3-Isopropyl-5-methylsulfanyl-7-(2-phenylbenzyl)amino-1(2)H-pyrazolo[4,3-d]pyrimidine (2)

A solution of 7-chloro-3-isopropyl-5-methylsulfanyl-1(2)H-pyrazolo[4,3-*d*]pyrimidine ***1*** [24] (700 mg, 2,88 mmol), 2-phenylbenzylamine (0.56 g, 3.15 mmol) and 0.4 mL trimethylamine in 10 mL acetonitrile was heated with stirring at 70 °C for 8 h. The solution was evaporated to dryness in vacuum, and the residue was purified by column chromatography on silica gel, stepwise with 2% and 4% MeOH in CHCl_3_. The product was obtained as white crystals from CHCl_3_ (763 mg, 68% yield), m.p. 82-86 °C, MS ESI^+^ 390.2 [M + H]^+^, ESI^−^ 388.1 [M – H]^-. 1^H (500 MHz; CDCl_3_): δ = 1.27 (d, *J* = 7.0 Hz, 6H, -CH(C*<u>H</u>*_3_)_2_); 1.43 (s, 3H, CH_3_); 2.39 (s, 3H, CH_3_); 3.20 (sept., *J* = 7.0 Hz, 1H, -C*<u>H</u>*(CH_3_)_2_); 7.72 (d, *J* = 5.2 Hz, 2H, NH-C*<u>H</u>*_2_-); 6.77 (s, 1H, N*<u>H</u>*-CH_2_-); 7.16-7.25 (m, 4H, H_Ar_); 7.28-7.33 (m, 4H, H_Ar_); 7.37 (d, *J* = 7.3 Hz, 1H, H_Ar_); ^13^C (125 MHz; CDCl_3_): δ = 14.3, 21.4, 26.0, 26.8, 42.2, 127.1, 127.6, 128.1, 128.5, 128.9, 130.0, 135.0, 140.4, 141.6, 163.2. Anal. Calcd. (C_22_H_23_N_5_S) C C, 67.84; H, 5.95 N, 17.98; S, 8.23. Found: C, 67.70; H, 6.06 N, 17.74; S, 8.15.

### 3-Isopropyl-7-(2-phenylbenzyl)amino-1(2)H-pyrazolo[4,3-d]pyrimidin-5-thiol (3)

3-Isopropyl-5-methylsulfanyl-7-(2-phenylbenzyl)amino-1(2)H-pyrazolo[4,3-*d*]pyrimidine **2** (0.383 g, 1 mmol) was dissolved in 2.65 mL of HMPA and sodium (56 mg, 2.5 eq.) was added under argon. The reaction mixture was stirred for 3 hours, and during this time, the temperature was continuously increased to 105 °C. For another 3 hours, the reaction mixture was maintained at 105 °C. The solution was cooled to 5 °C, and water (8 mL) was added. The solution was acidified by adding 5 N HCl. The precipitated crude product was isolated by filtration and purified by column chromatography on silica gel, stepwise with 1%, 2% and 3% MeOH in CHCl_3_. The product was obtained as yellow crystals (140 mg, 37% yield), m.p. 130-136 °C, MS ESI^+^ 376.1 [M + H]^+^, ESI^−^ 374.1 [M – H]^-^. NMR ^1^H (500 MHz; DMSO-*d*_6_): δ = 1.24 (d, *J* = 6.9 Hz, 6H, -CH(C*<u>H</u>*_3_)_2_); 3.40-3.47 (m, 1H, -C*<u>H</u>*(CH_3_)_2_); 4.64 (d, *J* = 5.5 Hz, 2H, NH-C*<u>H</u>*_2_-); 7.21-7.58 (m, 9H, H_Ar_); 9.02 (s, 1H, N*<u>H</u>*-). ^13^C (125 MHz; DMSO-*d*_6_): δ = 21.9, 24.0, 41.2, 127.1, 127.2, 127.6, 128.4, 129.2, 129.9, 135.5, 140.3, 141.0, 179.1. Anal. Calcd. (C_21_H_21_N_5_S) C, 67.17; H, 5.64 N, 18.65; S, 8.54, Found: C, 66.83; H, 5.61 N, 18.99; S, 8.85.

### 5-Piperidin-4-yl)thio-3-isopropyl-7-(2-phenylbenzyl)amino-1(2)H-pyrazolo[4,3-d]pyrimidine (4, LGR6768)

A solution of 3-isopropyl-7-(2-phenylbenzyl)amino-1(2)H-pyrazolo[4,3-*d*]pyrimidin-5-thiol **3** (98 mg, 0.26 mmol), 1-Boc-4-bromopiperidine (85 mg, 0.32 mmol) and KOH (25 mg, 0,45 mmol) in 2 mL of DMF was stirred for 12 h at room temperature. The solution was evaporated under vacuum, and the product was purified by column chromatography (stepwise 0.8%, 1.5% and 2% MeOH in CHCl_3_). The Bocylated product was dissolved in 0.5 mL MeOH, and then 0.5 ml of 10 N aq. HCl was added, and the solution was stirred for 1/2 h at 40 °C. The solution was evaporated under vacuum, and the rest was repeatedly evaporated with abs. EtOH until the product was obtained as an amorphous colourless glass foam (47 mg, 39% yield). MS ESI^+^ 459.2 [M + H]^+^, ESI^−^ 457.1 [M – H]^-^. 1H NMR (400 MHz, DMSO-*d*_6_) δ = 1.35 (d, *J* = 7.0 Hz, 6H, -CH-(C*<u>H</u>*_3_)_2_), 1.42 - 1.54 (m, 2H, -CH_2_-), 1.96 - 1.98 (m, 2H, -CH_2_-), 2.57 – 2.60 (m, 2H, -CH_2_-), 2.95 – 2.96 (m, 2H, -CH_2_-), 3.23 (sept., *J* = 6.9 Hz, 1H, -C*<u>H</u>*- (CH_3_)_2_), 3.68 (tt, *J* = 10.4, 3.5 Hz, 2H, -C*<u>H</u>*-S-), 4.58 (d, *J* = 4.3, 2H, Ar-C*<u>H</u>*_2_-NH-), 7.23 - 7.29 (m, 1H, Ar), 7.33 - 7.40 (m, 3H, Ar), 7.44 (d, *J* = 4.0, 5H, Ar), 8.60 (s, 1H, -NH). 13C NMR (100 MHz, DMSO-*d*_6_) δ = 21.7, 26.2, 31.6, 40.1, 41.4, 44.7, 127.2, 127.3, 127.6, 127.9, 128.3, 129.0, 129.8, 135.7, 138.8, 140.3, 141.0, 146.4, 149.8, 159.8. Anal. Calcd. (C_26_H_30_N_6_S. HCl) C, 63.08; H, 6.31; Cl, 7.16; N 16.97; S, 6.48, Found: C, 62.81; H, 6.67; N, 16.63; S, 6.23.

### 4.2. Crystallization, diffraction data collection, structure determination and analyses

CDK2/cyclin A2 protein was concentrated to 13.3 mg/mL in a buffer composed of 40 mM HEPES, 200 mM NaCl, and 0.02% monothioglycerol, pH 8.5, and mixed with 100 mM LGR6768 (in 100% DMSO). The final protein:inhibitor molar ratio was 1:3. Incubation was performed for 45 min on ice, and the sample was clarified by centrifugation at 4 °C (22,000×g, 15 min). Crystals were obtained at 18 °C using the sitting-drop vapour diffusion technique in Swissci 96-well 3-drop plates (Molecular Dimensions) and the Oryx8 robot (Douglas Instruments). The reservoir solution was created by mixing 23 µL of Morpheus (Molecular Dimensions) [33] condition #35 containing 10% w/v PEG 4,000, 20% v/v glycerol, 0.03 M NaNO_3_, 0.03 M Na_2_HPO_4_, 0.03 M (NH_4_)_2_SO_4_, 0.1 M Tris/Bicine pH 8.5 with 9 µL of JCSG^+^ (Qiagen) condition #29 containing 0.8 M sodium phosphate monobasic monohydrate, 0.8 M potassium phosphate monobasic, 0.1 M sodium HEPES, pH 7.5 [34]. Crystallization drops were prepared by mixing 100 nL of the protein-inhibitor complex and 200 nL of the reservoir solution. A plate-shaped crystal, size 150 µm × 100 µm × 10 µm, appeared after 3 days and was fished and flash-cooled in liquid N_2_ after 5 days.

A complete dataset at 2.6 Å resolution was collected at 100 K at beamline MX14.1 operated by Helmholtz-Zentrum Berlin (HZB) at the BESSY II electron storage ring (Berlin-Adlershof, Germany) [35]. The dataset was processed using the program XDS [36], and the structure was solved by the molecular replacement method using MolRep version 11.2.08 [37]with the CDK2/cyclin A2 complex available in the Protein Data Bank (PDB) under the code 6GVA [25] as the search model. Model refinement was performed using REFMAC 5.8.0352 from the CCP4 package version 8.0.002 [38] in combination with manual refinement in WinCoot version 0.9.8.1 [39]. The ligand model and restraints were prepared using the Ligand Builder feature in WinCoot [39]. Data collection and refinement statistics are shown in Table S1. The structural model was validated using MolProbity [40] and wwPDB [41] validation servers. Structural analysis was conducted using LigPlot+ [42] and the PISA server [43]. The figures were prepared using PyMol version 2.5.3 [44]. Atomic coordinates and structure factors were deposited into the PDB under the code 8B54.

### 4.3. Molecular modelling

Molecular modelling was performed on the cryo-EM structure of CDK7 (PDB: 7B5O). The 3D structure of LGR6768 was obtained from the cocrystal structure with CDK2/cyclin A2 (deposited as PDB: 8B54). Polar hydrogens were added to the ligand and protein with the AutoDock Tools program, and rigid docking was performed using AutoDock Vina 1.059 [45]. Interactions between LGR6768 and CDK7 were determined using BIOVIA Discovery Studio Visualizer 2021 (Dassault Systemes). Other figures were generated using PyMOL ver. 2.0.4 (Schrödinger, LLC).

### 4.4. Cell culture and viability assay

Human cell lines were obtained from the European Collection of Authenticated Cell Culture, German Collection of Microorganisms and Cell Cultures, Cell Line Service, and American Tissue Culture Collection or were kindly gifted by Jan Bouchal from Palacký University Olomouc (see Table S3 for more details and the cultivation conditions). The cells were grown in appropriate media supplemented with 10% foetal bovine serum, streptomycin (100 μg/ml), penicillin (100 IU/ml) and glutamine (4 mM) and stored in a humidified incubator at 37 °C and in 5% CO_2_.

For viability assays, the cells were seeded into 96-well plates and treated in triplicate with 6 different concentrations of LGR6768 for 72 hours. After the treatment, the resazurin solution (Sigma Aldrich) was added for 4 hours, and then the signal of resorufin was measured using a Fluoroskan Ascent microplate reader (Labsystems) at 544 nm/590 nm (excitation/emission). The GI_50_ (the drug concentration that is lethal for 50% of cells) was calculated from the dose response curves in Origin 6.0 software.

### 4.5. Kinase inhibition assay and selectivity profiling

The kinase inhibition assay was performed as previously described [46]. Briefly, all tested kinases were incubated with specific substrates in the presence of ATP, 0.05 µCi [γ^33^P]ATP and the LGR6768 compound in a final volume of 10 µl. The reaction was stopped by adding 3% aq. H_3_PO_4_. Aliquots were spotted onto P-81 phosphocellulose (Whatman), washed three times with 0.5% aq. H_3_PO_4_ and air-dried. Kinase inhibition was quantified using a FLA-7000 digital image analyser (Fujifilm). The drug doses required to decrease kinase activities by 50%, represented by the IC_50_ values, were determined from dose□response curves.

Kinase selectivity profiling was performed by Eurofins Discovery. The tested compound was assayed at a single concentration (1 µM) at ATP concentrations equal to K_m_.

### 4.6. Immunoblotting and antibodies

Cell lysates were separated by SDS□PAGE and electroblotted onto nitrocellulose membranes. The membranes were blocked, incubated with primary antibodies overnight, washed and incubated with secondary antibodies conjugated with peroxidase (Cell Signalling). Peroxidase activity was detected using Super-Signal West Pico reagents (Merck) using a CCD camera LAS-4000 (FujiFilm). The specific antibodies were purchased from Cell Signalling (anti-caspase 7; anti-caspase 9; anti-cdc2; anti-CDK2, clone 78B2; anti-cyclin A2, clone BF683; anti-cyclin B1, clone V152; anti-HSP60, clone D6F1; anti-Mcl-1, clone D35A5; anti-PARP, clone 46D11), Santa Cruz Biotechnology (anti-β-actin, clone C4; anti-caspase-3; anti-p-Cdc2 p34 (Thr 161); anti-p-CDK2 (Thr 160)) or Merck (anti-Bcl-2, clone Bcl-2-100; anti-α-tubulin, clone DM1A).

### 4.7. Caspase 3/7 activity assay

Cell lysates were incubated with 100 µM Ac-DEVD-AMC (Enzo Life Sciences), which is a substrate for caspases 3 and 7, in assay buffer (25 nM PIPES, 2 mM MgCl_2_, 2 mM EGTA, 5 mM DTT, pH 7.3). The fluorescent signal of the product was measured at 355 nm/460 nm (excitation/emission) using a Fluoroskan Ascent microplate reader (Labsystems).

### 4.8. Cell cycle analysis

After harvesting, the cells were then washed with PBS, fixed with 70% ethanol, denatured with 2 M HCl and neutralized. After staining with propidium iodide, the cells were analysed by flow cytometry using a 488 nm laser (BD FACS Verse with BD FACSuite software, version 1.0.6.). Cell cycle distribution was analysed using ModFit LT (Verity Software House, version 4.1.7).

### 4.9. RNA isolation and qPCR

Total RNA was isolated via an RNeasy plus mini kit (QIAGENE) according to the manufacturer’s instructions. The concentration and purity of isolated RNA were measured on a DeNovix DS-11 spectrophotometer. Isolated RNA (0.5-1 µg) was then used for reverse transcription into first strand cDNA via a SensiFast cDNA synthesis kit (Bioline). RNA spike I (TATAA) was used as an inhibition control of the transcription reaction. Quantitative PCR was carried out with a SensiFAST SYBR No-Rox Kit (Bioline) on a CFX96 Real-time PCR Detection System. Suitable primers were designed via Primer BLAST [47] and synthesized by Generi Biotech. Raw data were analysed using CFX Maestro 2.2, and relative normalized expression was commutated by the ΔΔC_T_ method [48]. Expression was normalized to GAPDH and RPL13A in the MOLM-13 cell line or GAPDH and B2M in the SEM cell line.

Those pairs of genes were found to be the most stable for concrete cell lines using CFX Maestro 2.2 software.

List of used primers:

BCL-2: F – ATGTGTGTGGAGAGCGTCA, R – ACAGCCAGGAGAAATCAAACAG

GAPDH: F – TCCAAAATCAAGTGGGGCGA, R – TGGTTCACACCCATGACGAA

MCL-1: F-AGTTGTACCGGCAGTCG, R – TTTGATGTCCAGTTTCCGAAG

MYB: F – TCTCCAGTCATGTTCCATACCC, R – TGTGTGGTTCTGTGTTGGTAGC

MYC: F – TACAACACCCGAGCAAGGAC, R – AGCTAACGTTGAGGGGCATC

RPL13A: F – CGACAAGAAAAAGCGGATGG, R -TTCTCTTTCCTCTTCTCCTCC

RUNX1: F-CACTGTGATGGCTGGCAATG, R: CTTGCGGTGGGTTTGTGAAG

RUNX2: F – TCTGGCCTTCCACTCTCAGT, R: TGGCAGGTAGGTGTGGTAGT

HOXA9: F – CCTGACTGACTATGCTTGTGGT, R -ACTCTTTCTCCAGTTCCAGGG

## Acknowledgements

Authors would like to thank Veronika Vojáčková for excellent technical support. The work was supported by the European Union -Next Generation EU (The project National Institute for Cancer Research, Programme EXCELES, Project No. LX22NPO5102), Czech Science Foundation (21-06553S) and Palacký University Olomouc (IGA_PRF_2022_007).

