## Supplementary Material for "Characterization of new highly selective pyrazolo[4,3-d]pyrimidine inhibitor of CDK7"

**Content**

1. **Synthesis of LGR6768 and crystal structure description**

   Figure S1-S3, Table S1

2. **Kinase selectivity**

   Figure S4, Table S2

3. **Further cellular tests**

   Figure S5

4. **NMR spectra**

5. **Cultivation condition of used cell lines**

   Table S3

**Synthesis of LGR6768 and crystal structure description**

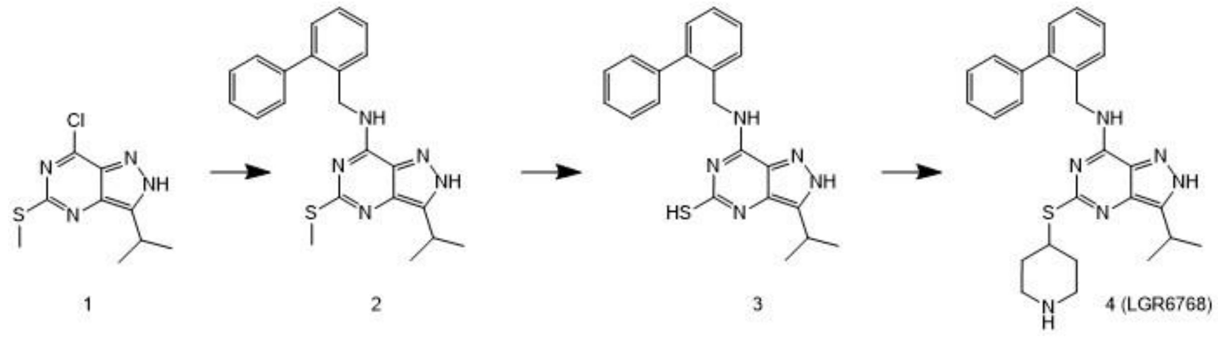


**Figure S1:** Synthesis of LGR6768

**Table S1:** X-ray data collection and refinement statistics

| **Data collection statistics** | |
|---|---|
| Space group | $P2_12_12_1$ |
| Cell parameters a, b, c [Å]; a, β, γ [º] | 73.02, 134.99, 163.96<br>90.00, 90.00, 90.00 |
| Wavelength [Å] | 0.918 |
| Resolution [Å] | 50.00-2.60 (2.76-2.60) |
| Unique reflections | 50,620 (8,027) |
| Multiplicity | 13.6 (14.2) |
| Completeness [%] | 99.9 (100) |
| $R_{meas}$ [%][a] | 61.3 (290.3) |
| $CC_{(1/2)}$ [%][b] | 98.8 (71.7) |
| Average I/σ(I) | 5.42 (1.01) |
| Wilson B [$Å^2$][c] | 39.44 |
| **Refinement statistics** | |
| Resolution range [Å] | 49.56-2.60 (2.68-2.60) |
| No. of reflections in working set | 49,156 (3,539) |
| No. of reflections in test set | 1,460 (105) |
| R value [%][d] | 23.3 (40.3) |
| R-free value [%][e] | 28.7 (39.8) |
| RMSD deviation from ideal bond length [Å] | 0.011 |
| RMSD deviation from ideal bond angle [°] | 1.596 |
| Number of protein atoms | 8,845 |
| Number of water molecules | 314 |
| Number of other non-protein atoms | 98 |
| Mean B value [$Å^2$] | 39.61 |
| Residues in Ramachandran favored regions [%][f] | 96.47 |
| Residues in Ramachandran allowed regions [%][f] | 99.63 |

[a]$R_{meas}$ defined in ref. [1]. [b]$CC_{(1/2)}$ is Pearson's correlation coefficient determined on the data set randomly split in half [2]. [c]Wilson B by the Sfcheck program from the CCP4 suite [3]. [d]R-value = $||F_o| - |F_c||/|F_o|$, where $F_o$ and $F_c$ are the observed and calculated structure factors, respectively. [e]$R_{free}$ is

equivalent to the R-value but is calculated for 5% of the reflections chosen at random and omitted from the refinement process [4]. [f]As determined by MolProbity [5].

The asymmetric unit of the crystal contains two CDK2/cyclin A2 heterodimers (protein chains A/B and C/D). All the residues are modeled into the electron density map, except for disordered residues 39-40 in chains A and C, as well as residues 175-176 in chain B and 175-177 in chain D. Inhibitor LGR6768 was modeled into well-defined electron density in both active sites with full occupancy (Figure S2). The RMSD for the superposition of the Cα atoms for the two heterodimers is 0.154 $Å^2$ and the conformation of the active site residues and the ligand poses in the two heterodimers are almost identical. Heterodimer A/B was used to describe protein – inhibitor interactions.

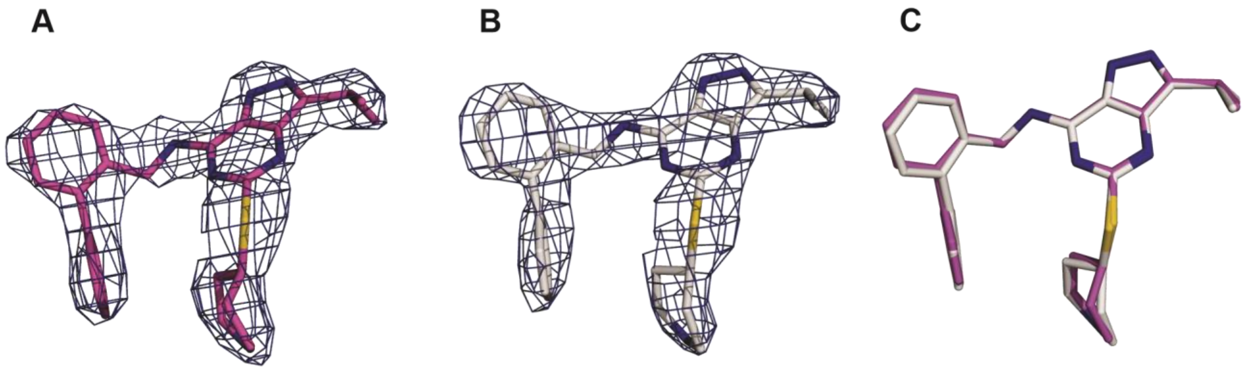


**Figure S2:** Electron density for the inhibitor LGR6768 in the active site A (panel A) and C (panel B). A superposition of the inhibitor molecules bound in the two active sites is shown in panel C. The inhibitor is shown as sticks with carbon atoms in magenta (active site A) and gray (active site C), nitrogen atoms are blue, and sulfur atoms yellow; *2Fo-Fc* maps are shown as blue mesh at 1.3 σ.

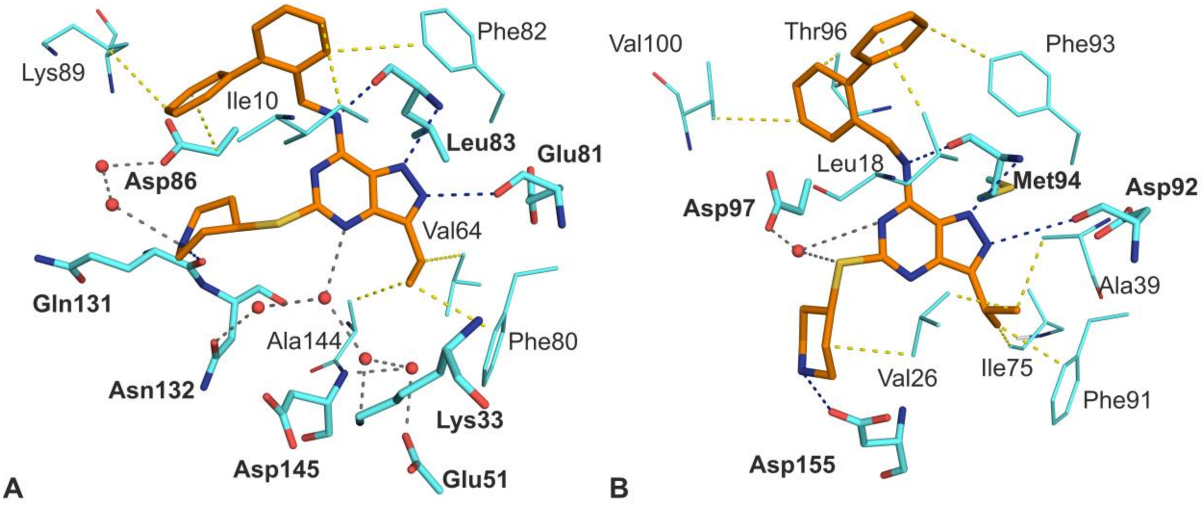


**Figure S3:** LGR6768 in the active site of CDK2 (crystal structure, panel A) and CDK7 (molecular docking, panel B). Interacting residues are shown as sticks with H-bond forming residues labelled with bold font and other residues with regular font. Direct H-bonds are shown as blue dashed lines, water-mediated H-bonds as grey dashed lines, and hydrophobic interactions as yellow dashed lines. Carbon atoms are cyan (protein) and orange (inhibitor). Nitrogen atoms are blue, oxygen atoms red, sulphur atoms are yellow, and waters are shown as red spheres. Residues Ala31 Val18, and Leu134 from CDK2, and Leu144 from CDK7 were not shown for clarity. Structure of LGR6768 in complex with CDK2 has been deposited to the PDB under the code 8B54.

#### Kinase selectivity

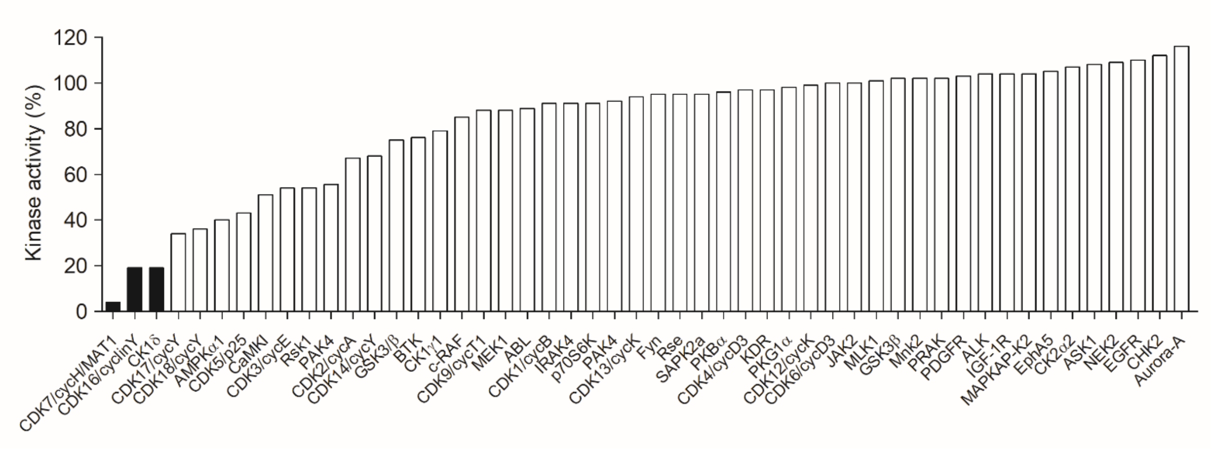


**Figure S4:** Selectivity profiling of LGR6768. The compound was applied in a single 1 μM concentration on 50 kinases.

**Table S2:** CDK selectivity profile of LGR6768

| kinase | $IC_{50}$ (μM)* |
|---|---|
| **CDK1/cyclin B** | 5.14 (± 0.97) |
| **CDK2/cyclin A** | 0.50 (± 0.05) |
| **CDK2/cyclin E** | 0.25 (± 0.01) |
| **CDK4/cyclin D1** | >20 |
| **CDK5/p25** | 1.12 (± 0.1) |
| **CDK7/cyclin H/MAT1** | 0.02 (± 0.02) |
| **CDK9/cyclin T** | 6.55 (± 1.70) |

*** – measured at least in duplicate**

**Further cellular tests**

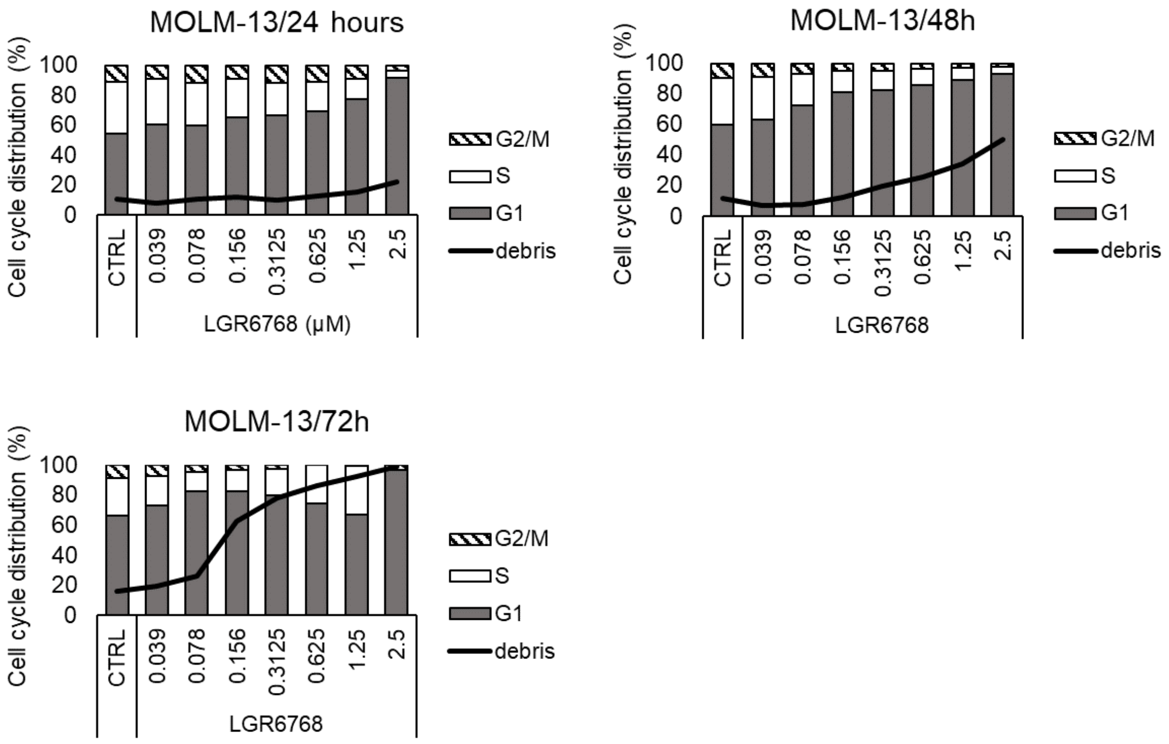


**Figure S5:** Effect of LGR6768 on cell cycle in MOLM-13 cell line. Cells were treated for 24, 48, or 72 hours with an increasing concentration of LGR6768 or vehicle.

### NMR Spectra

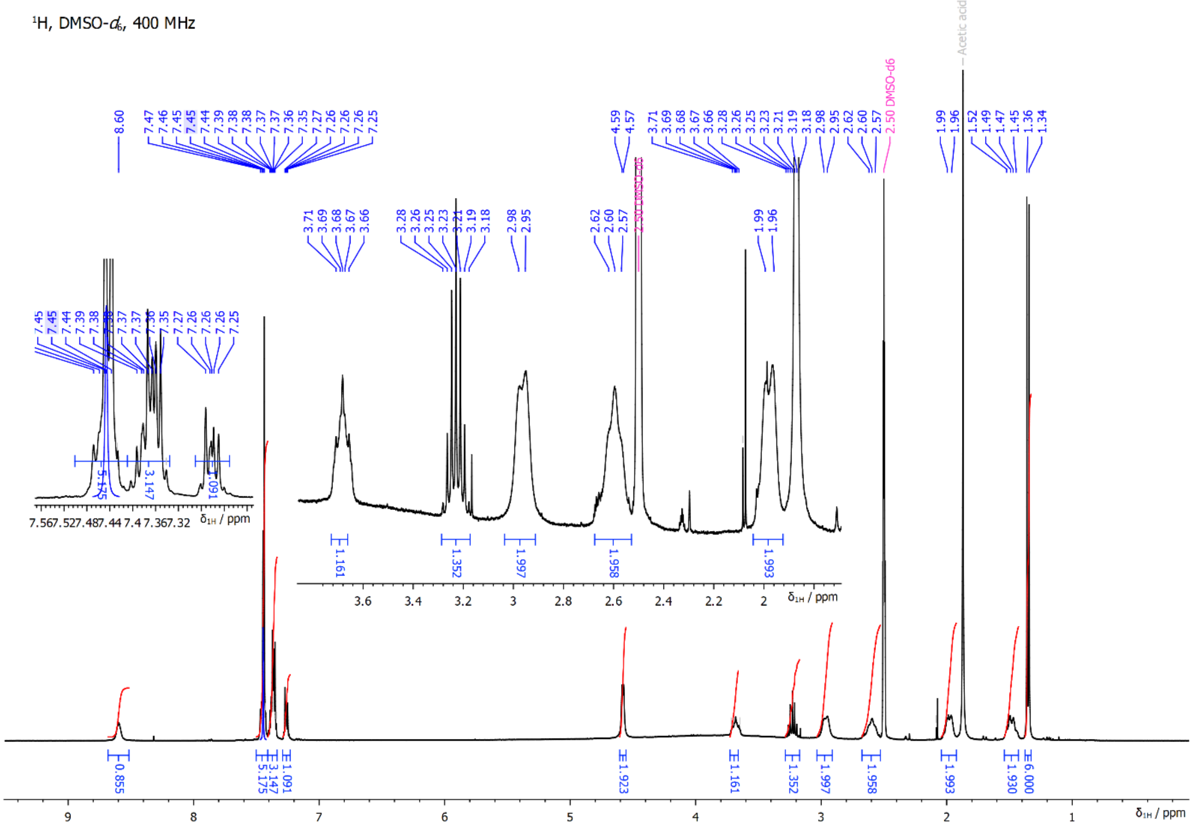


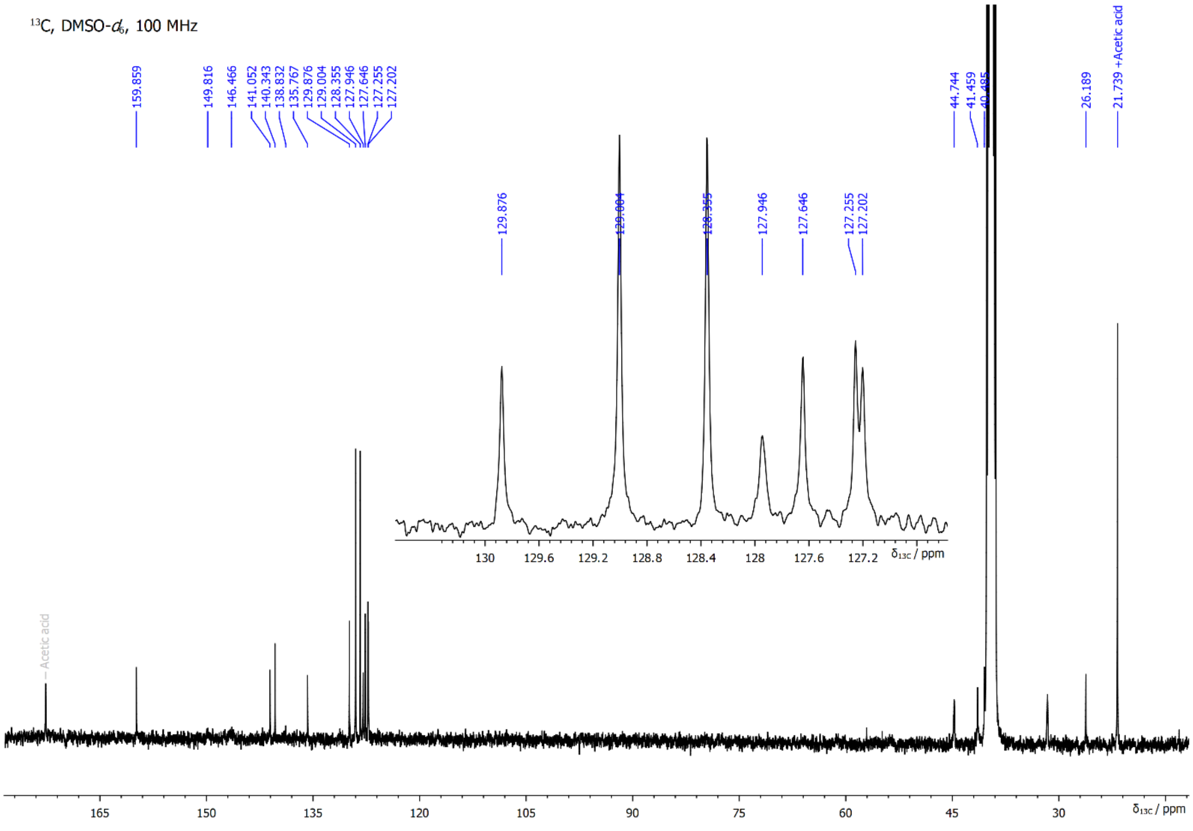

**Cultivation conditions of used cell lines**

**Table S3:** Used cell lines with source and culture media

| Cell line | Cancer type | Source | Used medium |
|---|---|---|---|
| **22Rv1** | Prostate | Jan Bouchal* | RPMI |
| **BT474** | Breast | Jan Bouchal* | DMEM |
| **CCFR- CEM** | ALL | ECACC | RPMI |
| **DU145** | Prostate | Jan Bouchal* | RPMI |
| **G361** | Melanoma | ECACC | DMEM |
| **HL-60** | AML | ECACC | IMDM |
| **K562** | CML | ECACC | DMEM |
| **LAPC4** | Prostate | Jan Bouchal* | DMEM |
| **MCF7** | Breast | ECACC | DMEM |
| **MINO** | Lymphoma | ATCC | RPMI |
| **MOLM-13** | AML | DSMZ | RPMI |
| **MV4-11** | AML | Cell line service | RPMI |
| **RS4-11** | ALL | DSMZ | α-MEM |
| **SEM** | ALL | DSMZ | IMDM |
| **SKBR3** | Breast | ATCC | DMEM |
| **THP-1** | AML | DSMZ | RPMI |
| **U937** | AML | DSMZ | RPMI |

AML – acute myeloid leukemia, ALL – acute lymphoblastic leukemia, CML – chronic myeloid leukemia; ATCC **-** American Tissue Culture Collection; DSMZ **-** German Collection of Microorganisms and Cell Cultures; ECACC - European Collection of Authenticated Cell Culture;

***** Cells were kindly gifted by Jan Bouchal from Palacký University Olomouc.
